# CPO Complete, a novel test for fast, accurate phenotypic detection and classification of carbapenemases

**DOI:** 10.1101/710939

**Authors:** Gina K. Thomson, Sameh AbdelGhani, Kenneth S. Thomson

**Author notes:** Corresponding Author: (KST). These authors contributed equally to this work (Gina K. Thomson, Kenneth S. Thomson). This author also contributed to this work (Sameh AbdelGhani).

## Abstract

Rapid, accurate detection of carbapenemase-producing organisms (CPOs) and the classification of their carbapenemases are valuable tools for reducing the mortality of the CPO-associated infections, preventing the spread of CPOs, and optimizing use of new β-lactamase inhibitor combinations such as ceftazidime/avibactam and meropenem/vaborbactam. The current study evaluated the performance of CPO Complete, a novel, manual, phenotypic carbapenemase detection and classification test. The test was evaluated for sensitivity and specificity against 262 CPO isolates of *Enterobacteriaceae, Pseudomonas aeruginosa* and *Acinetobacter baumannii* and 67 non-CPO isolates. It was also evaluated for carbapenemase classification accuracy against 205 CPOs that produced a single carbapenemase class. The test exhibited 100% sensitivity 98.5% specificity for carbapenemase detection within 90 minutes and detected 74.1% of carbapenemases within 10 minutes. In the classification evaluation, 99.0% of carbapenemases were correctly classified. The test is technically simple and has potential for adaptation to automated instruments. With lyophilized kit storage at temperatures up to 38°C the CPO Complete test has the potential to provide rapid, accurate carbapenemase detection and classification in both limited resource and technologically advanced laboratories.

## Background

Infections by carbapenemase-producing organisms (CPOs) are associated with high mortality (1-4). The main contributors to this are the extensive, sometimes total, antibiotic resistance of CPOs, failures to provide effective therapy, and inadequate infection control. Rapid CPO detection and carbapenemase classification can be pivotal to efforts to reduce the mortality. Speed of detection is critical for effective infection control and for prompt initiation of combination antibiotic therapy, which is optimal for serious CPO infections (2, 3, 5-16). Classification of carbapenemases can optimize appropriate use of new β-lactamase inhibitor combinations such as ceftazidime/avibactam, meropenem/vaborbactam, imipenem/relebactam, aztreonam/avibactam, meropenem/nacubactam, cefepime/zidebactam and cefepime/VNRX-5133 and minimize their overuse. Currently marketed rapid, phenotypic manual CPO tests do not classify carbapenemases.

The CPO Complete test is a manual CPO detection and classification test designed to provide rapid, accurate results. The current study was designed to assess its speed and accuracy of CPO detection and its potential to classify carbapenemases.

## Materials and Methods

### Isolates

Three hundred twenty nine isolates of *Enterobacteriaceae, Pseudomonas aeruginosa* and *Acinetobacter baumannii* from two culture collections, that of the University of Louisville and also the Antimicrobial Resistance Isolate Bank of the Centers for Disease Control and Prevention and Food and Drug Administration, were characterized for types of β-lactamase production by PCR, microarray, DNA sequencing, whole genome sequencing, phenotypic and biochemical tests. Those tested in the carbapenemase detection phase of the study included isolates of high diagnostic difficulty. They included 125 isolates producing KPC, NMC-A, IMI, and SME class A carbapenemases; 87 isolates producing NDM, SPM, IMP, VIM and GIM class B carbapenemases; 44 isolates producing OXA-23, 40, 48, 58, 72, 181, and 232 class D carbapenemases; 6 isolates producing 2 carbapenemase classes; and 67 carbapenemase-negative isolates (non-CPOs) that included porin mutants and producers of ESBLs, AmpCs, K1, and broad spectrum β-lactamases. Classification potential was assessed for 205 CPOs that produced a single carbapenemase class. These comprised 185 isolates from the detection part of the study (88 class A, 56 class B, 41 class D) and an additional 10 class A (9 KPC, 1 SME), 7 class B (4 VIM, 3 NDM) and 3 OXA-48-like class D producers. Control strains were *K. pneumoniae* BAA 1705 (KPC), *E. coli* BAA 2452 (NDM-1), *A. baumannii* CDC 0035 (OXA-72), and *K. aerogenes* (formerly *E. aerogenes*) G1614 (non-CPO).

### Carbapenemase detection

The CPO Complete test did not require an initial lysis step prior to incubating the test. The test solution (solution A) was prepared by dissolving 12 mg of pharmaceutical imipenem/cilastatin (Hospira cat. no. NDC 0409-3507-21), 10 mg thimerosal (Enzo cat. no. ALX-400-013-G005), 5 mg glucose (Sigma cat no. G-5000) and 4 mg polymyxin B (EMD Millipore Corp., USA, cat. no. 5291-500MG) in a mixture of 1 ml of Mueller-Hinton broth (BD Diagnostics Systems, Sparks, MD), 30 μl zinc sulfate (Sigma-Aldrich Co. cat. no Z2876) and 140 μl phenol red solution (VWR International catalog # 97062-476). This solution was pH adjusted to pH 7.0 using 10N NaOH & 12N HCl.

Thirty μl of solution A was dispensed into transparent vessels such as PCR tubes (VWR International catalog # 20170-004) or microtiter wells. One tube was used for each test isolate and two additional tubes were used for testing a positive and negative control isolates. Colonies of each bacterial test isolate and a positive and negative control isolate were harvested with a 1 μl loop (VWR International, catalog # 12000-806) from blood agar (BD Diagnostics Systems, Sparks, MD, USA). The volume of harvested inoculum was sufficient to provide a slightly convex surface after filling the loop aperture e.g. similar to the amount of convex curvature of these two parentheses (). Excess inoculum was avoided to ensure test accuracy. The inoculum for each isolate was suspended in 30 μl of Solution A by vigorously rotating the loop in the solution. Inoculated tubes were then incubated at room temperature until positive or for a maximum of 90 minutes. The test was interpreted in bright light against a white background by comparing the color of the inoculated test to the negative control *K. aerogenes* G1614. A positive test was interpreted as the development of yellow, orange or a lighter shade of red than the red negative control (Fig 1). Tests were performed blinded.

**Fig 1.**
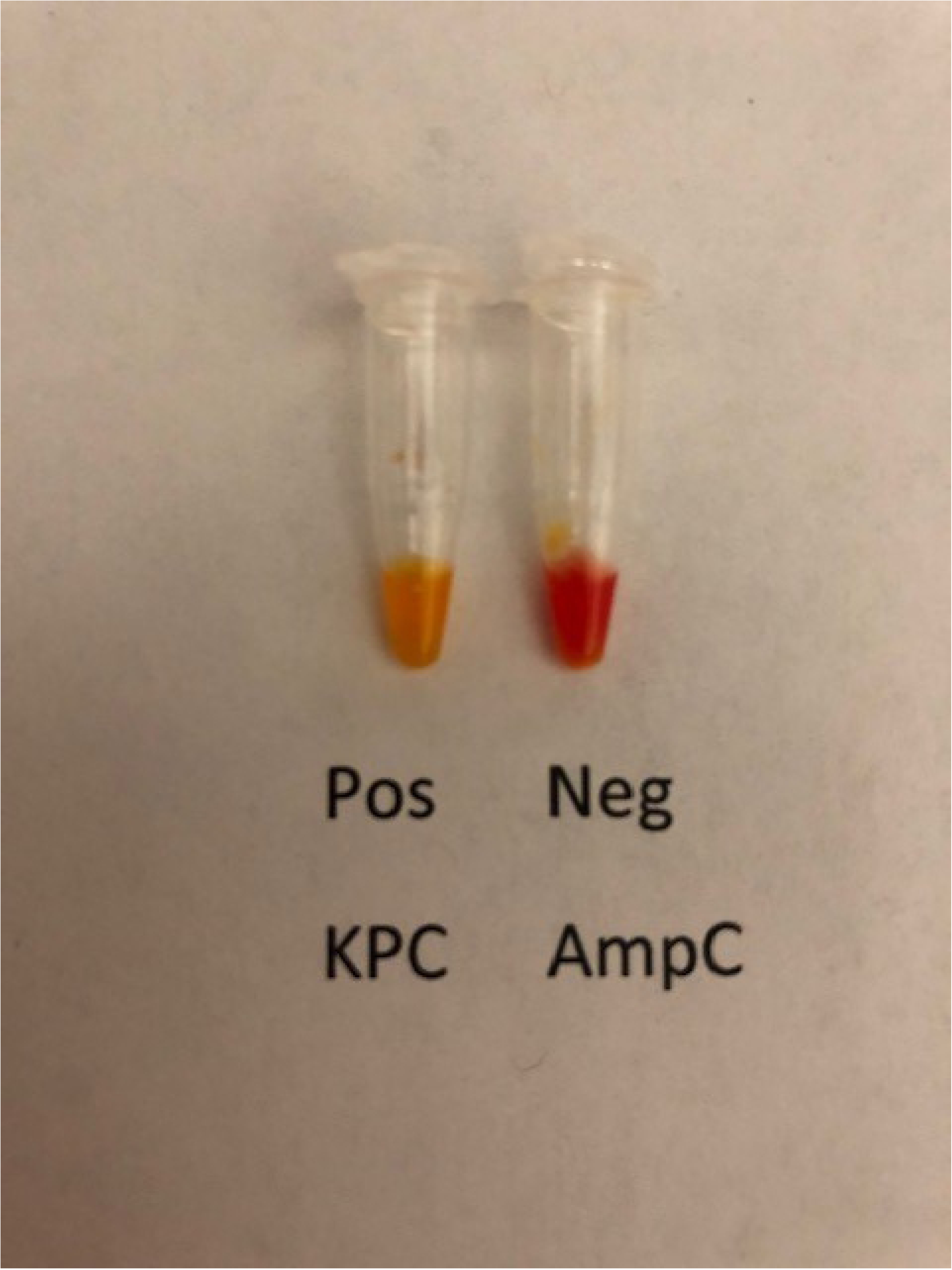
A positive test was interpreted as the development of yellow, orange or a lighter shade of red than the negative control. Tests with the positive control isolate, KPC-producing *K. pneumoniae* BAA 1705 and the negative control isolate *K. aerogenes* G1614 are shown.

### Carbapenemase classification

In contrast to the carbapenemase detection tests in which tests were performed blinded, the classification phase of the study was proof-of-concept testing with carbapenemase classifications known at the time of testing. Three solutions were used for classification. These were solutions A (described above), B and C. Solution B comprised 30 μl of solution A supplemented with 2 μl of a solution prepared by dissolving 120 mg of phenyl boronic acid in 3 ml dimethyl sulfoxide (VWR International catalog # BDH1115-1LP) and adding this to 3 ml sterile inoculum water (Beckman Coulter, Inc. Brea, CA, cat no B1015-2) and 840 μl phenol red solution. The final solution was pH adjusted to pH 7.5 using 10N NaOH and 12 N HCl. Solution C comprised 30 μl of solution A supplemented with 2 μl of a solution prepared by dissolving 235 mg pyridine-2,6-dicarboxylic acid, 98% (also known as dipicolinic acid) (Alfa Aesar cat no A12263) in 10 ml Tris-EDTA buffer solution 100x concentrate (Sigma T9285) and 1,400 μl phenol red solution. The final solution was pH adjusted to pH 6.8 using 10 N NaOH and 12N HCl.

Using a separate 1 μl loop for each tube, sets of the three solutions were inoculated for each of 205 isolates that produced a single carbapenemase. Tests with solutions B and C were inoculated by the same procedure used for tests with solution A. The isolates comprised 98 producers of class A carbapenemases, 63 producers of class B carbapenemases and 44 producers of class D carbapenemases. Only CPOs (i.e. positive result with solution A) were assessed for carbapenemase classification. Tests with solutions B and C scored as positive or negative by interpreting visually for a change from the initial red color to a lighter color. The interpretation was based on which of solutions B or C was more positive (i.e. lighter in color). Occasionally the color changes in solutions B or C were slower and less intense than the color change in solution A. Tests with solutions B and C were ignored if solution A yielded a negative result. Classifications were interpreted according to Table 1. Figs 2-4 show representative classification test results.

**Table 1.** Guide to Interpretation of Carbapenemase Classification Tests.

| Carbapenemase<br>Classification | Solution |  |  |
| --- | --- | --- | --- |
|  | A | B | C |
| **Class A** | Positive | Negative | Positive |
| **Class B** | Positive | Positive | Negative |
| **Class D** | Positive | Positive | Positive |
| **Positive Untyped** | Positive | Negative | Negative |
| **Negative** | Negative | Not applicable | Not applicable |

**Fig 2.**
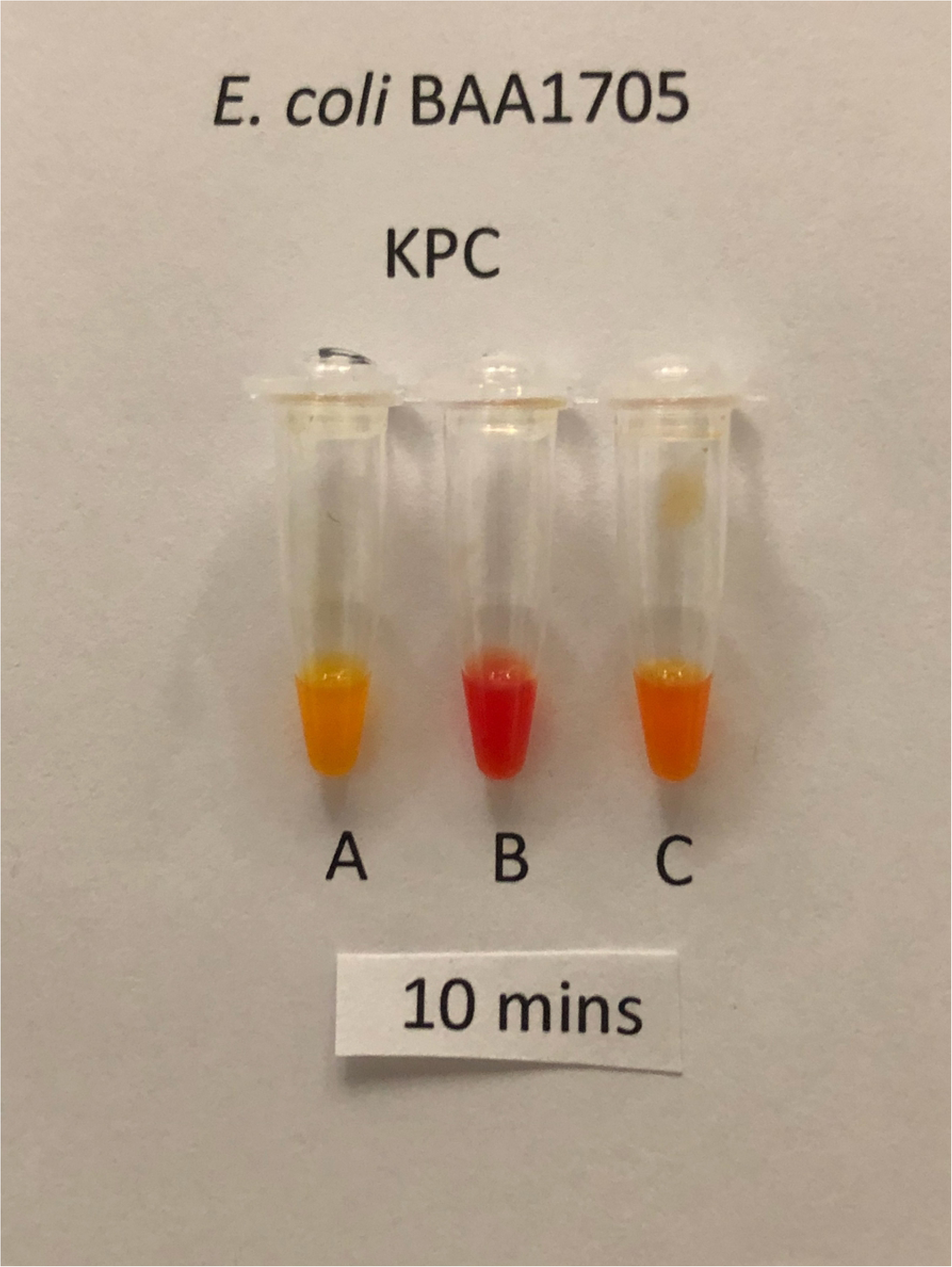
Classification test for Class A CPO, KPC-producing *K. pneumoniae* BAA 1705 after 10 minutes incubation. Tubes A and C are positive (yellow). Tube B is negative (red). Tube A contains only the test solution and indicates that the isolate is carbapenemase-positive. Tubes B and C contain the test solution plus inhibitors of Class A and B carbapenemases respectively.

**Fig 3.**
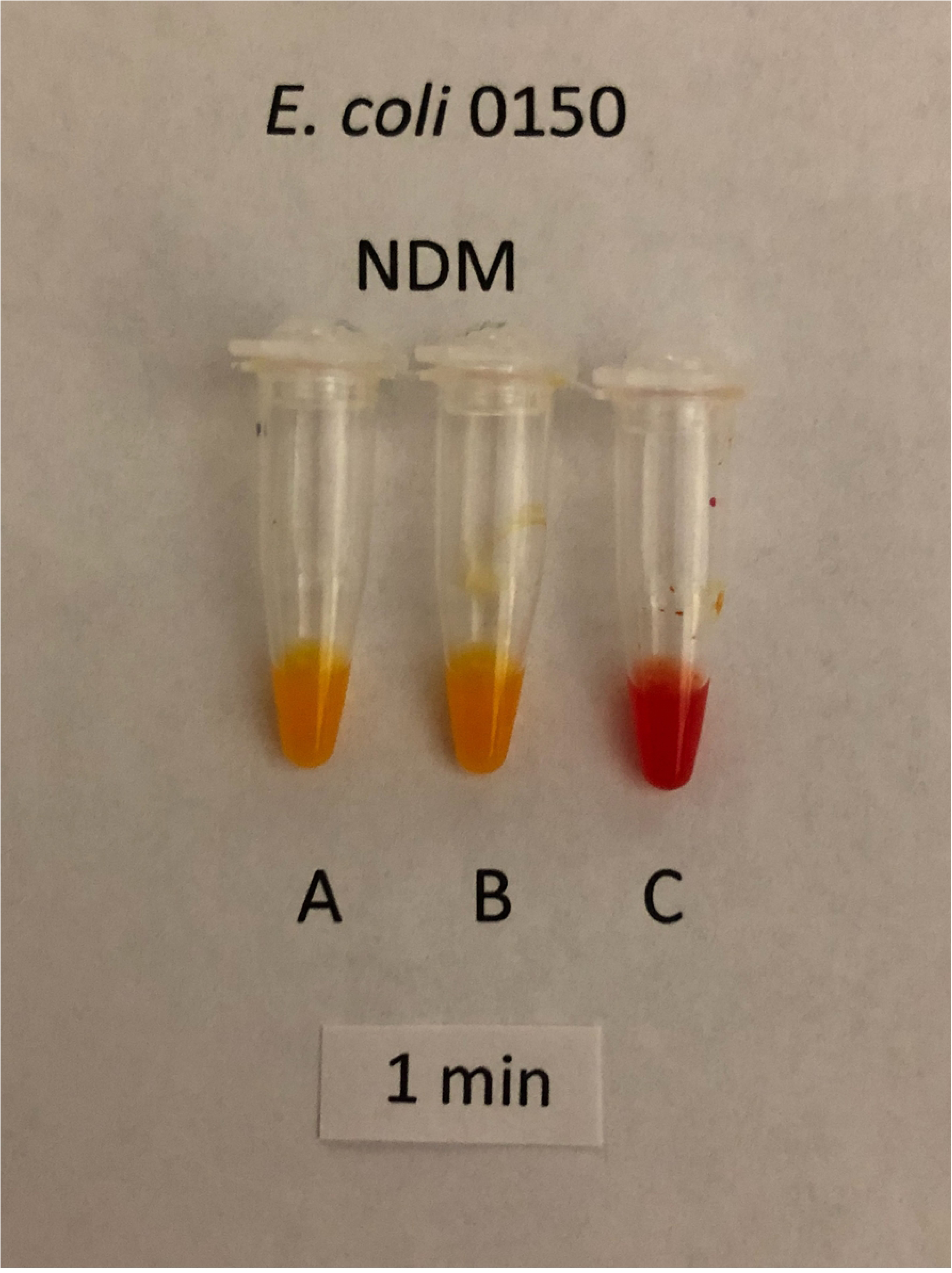
Classification test for Class B CPO, NDM-5-producing *E. coli* CDC 0150 after 1 minute incubation showing positive results in tubes A and B and a negative result in tube C.

**Fig 4.**
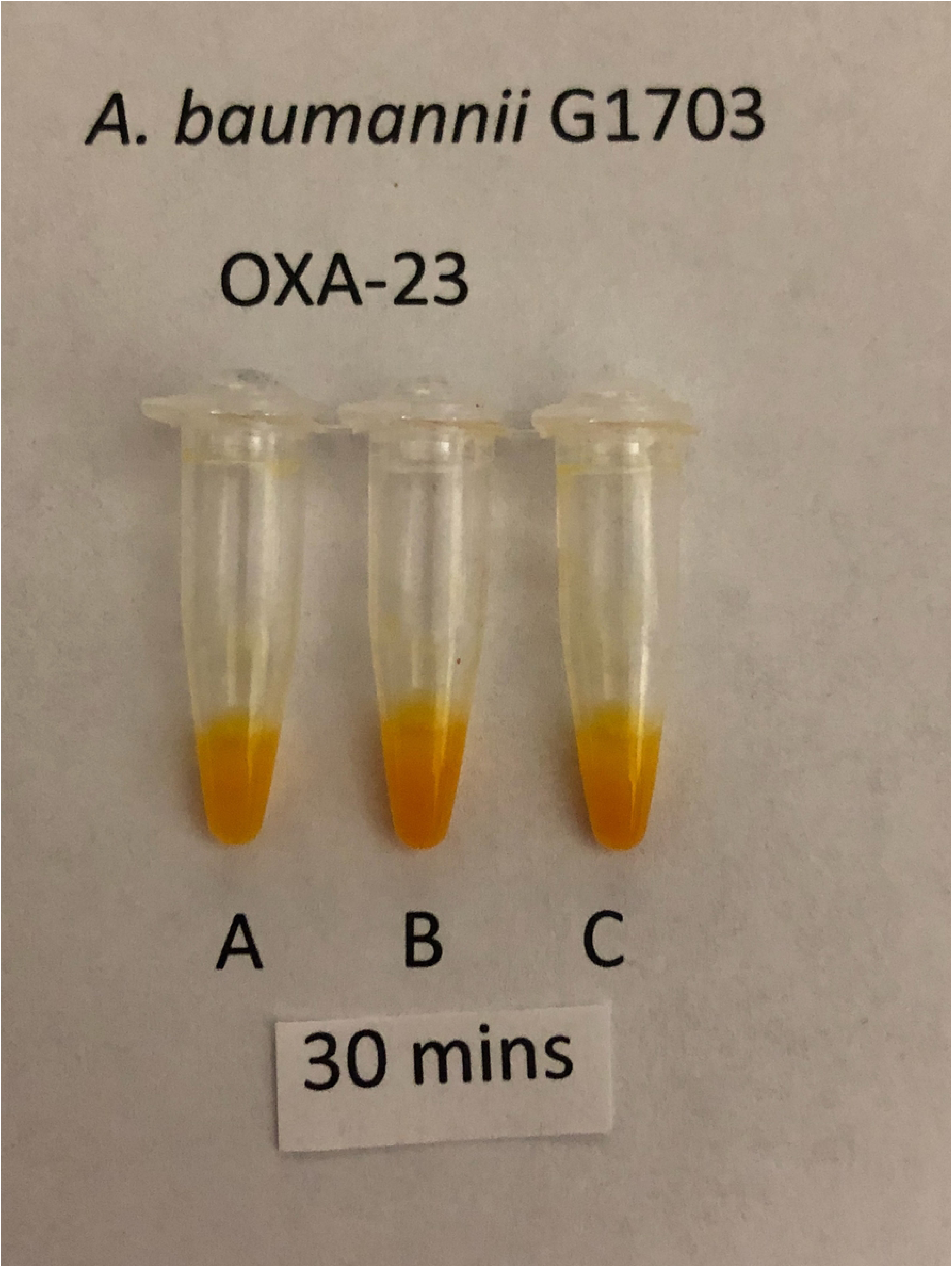
Classification test for Class D CPO, OXA-23-producing *A. baumannii* G1703 after 30 minutes incubation showing positive results in all three tubes.

## Results

### Carbapenemase Detection

All 262 CPOs yielded positive tests within 90 minutes (100% sensitivity) and only one of 67 non-CPOs yielded a falsely positive result (98.5% specificity) (Table 2). The carbapenemases of 74.1% of the CPOs were detected within seconds to 10 minutes and 98.1% were detected within 1 hour. Notably, KPC-producing *P. aeruginosa* isolates typically yielded positive results within 1 minute, 91.3% of isolates producing an OXA carbapenemase were positive within 60 minutes, and positive results were obtained with isolates of high diagnostic difficulty such as KPC-producing *A. baumannii*, KPC-4-producing *K. pneumoniae* and IMP-27-producing *Proteus mirabilis*. The single falsely positive result occurred with an AmpC-producing *E. cloacae* isolate.

**Table 2.**
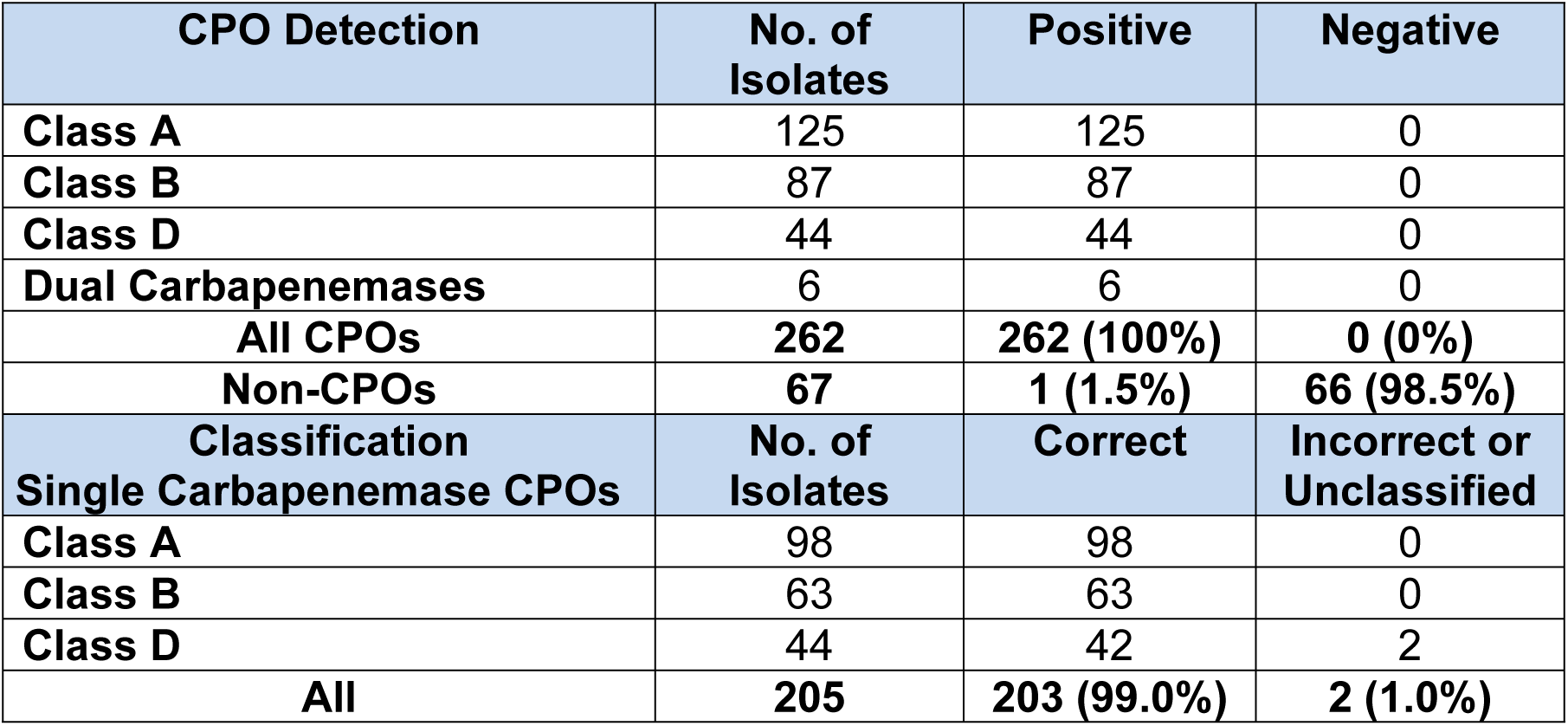
CPO Detection and Classification Test Results.

### Carbapenemase Classification

As shown in Table 2, 99% of carbapenemases were correctly classified. All class A and B carbapenemases were classified correctly and 42 of the 44 class D carbapenemases were classified correctly. Two class D carbapenemase producers yielded a positive but unclassified result.

## Discussion

The high mortality and continuing emergence of resistance of CPOs make it essential that laboratories provide a strong diagnostic contribution in meeting the need to reduce the mortality and control the spread of these pathogens. Rapid, accurate CPO detection can facilitate prompt, appropriate, targeted therapy and effective infection control measures (17). Detection tests that require overnight incubation are too slow for therapeutic and infection control needs but may be suitable for epidemiological studies (18, 19). It is also important to use detection tests that minimize falsely positive results as these can have adverse consequences for patients such as being repeatedly subjected to unwarranted infection control precautions and receiving suboptimal, toxic therapy such as polymyxins (20).

It is now vitally important that laboratories rapidly classify carbapenemases to inform clinicians about the potential therapeutic utility of the new β-lactamase inhibitor combinations. Identification of class A carbapenemases indicates that the currently FDA-approved agents, ceftazidime/avibactam and meropenem/vaborbactam, are potential therapeutic options, while the detection of a class B carbapenemase contraindicates these agents. Classification can be a clinically powerful source of information about whether or not these agents are contenders for therapy but it does not eliminate the need for antibiotic susceptibility testing (15).

Carbapenemase classification has a second vital role in antibiotic stewardship for prevention of emergence of resistance to the new β-lactamase inhibitor combinations. These agents provide the opportunity to safely treat patients with CPO infections and avoid highly toxic agents such as the polymyxins and aminoglycosides. It is of critical importance to ensure that the currently approved new β-lactamase inhibitor combinations are not overused and select resistance, not only to themselves, but possibly also to the other β-lactamase inhibitor combinations currently in development. In particular, the potential for development of resistance to avibactam (21, 22) is a concern as resistance to it may impart cross-resistance to the related diazobicyclooctane analogs, relebactam and nacubactam. Similarly, resistance to vaborbactam may impart cross-resistance to its chemically related counterpart, VNRX-5133. Carbapenemase classification is a tool that can help to prevent overuse and guide appropriate use of the current new inhibitor combinations so that they are used almost solely for infections caused by class A CPOs and are restricted for infections by pathogens with other resistance mechanisms. This is vital because, despite the efficacy of these agents for class A CPO infections (23-27), gram-negative resistance continues to emerge (18, 28-33) and threatens our current window of opportunity for treating CPO infections with agents less toxic than those previously available.

A strength of this study was the variety of genotypes and phenotypes tested that included isolates of considerable diagnostic difficulty. A limitation was the non-blinded nature of the carbapenemase classification testing. It is now necessary to determine the accuracy of classifications in blinded testing. It is also necessary to assess classification testing with isolates that produce more than one class of carbapenemase. It was, nevertheless, a promising finding that CPO Complete correctly classified 99% of carbapenemases and that no class B CPOs were misclassified as class A.

In conclusion, in this study the CPO Complete test detected all carbapenemases rapidly, with a 10-minute detection rate of 74.1%. The speed and accuracy of the test, coupled with its potential to classify carbapenemases, can be applied successfully to meeting what has become one of the world’s most urgent infectious disease challenges (3, 7, 13, 19, 34-42). In addition to its accurate performance, the test has potential for incorporation in currently available automated susceptibility tests. Unlike molecular probes that may be able to detect only a small number of molecular targets and unable to distinguish between carbapenemase and non-carbapenemase *bla*_KPC_ variants, this and other phenotypic tests are likely to become increasingly useful as KPC variants continue to emerge (17, 33).

Furthermore, CPO Complete can be stored lyophilized at temperatures up to 38°C making it amenable to implementation in both limited resource and technologically advanced laboratories. In all, these features suggest that CPO Complete can contribute to the challenges of improving patient management, reducing CPO-associated mortality, and containing the spread of CPOs.

